# Metavisitor, a suite of Galaxy tools for simple and rapid detection and discovery of viruses in deep sequence data

**DOI:** 10.1101/048983

**Authors:** Guillaume Carissimo, Marius van den Beek, Juliana Pegoraro, Kenneth D Vernick, Christophe Antoniewski

**Author notes:** These authors contributed equally to the work (Guillaume Carissimo, Marius van den Beek).

## Abstract

We present user-friendly and adaptable software to provide biologists, clinical researchers and possibly diagnostic clinicians with the ability to robustly detect and reconstruct viral genomes from complex deep sequence datasets. A set of modular bioinformatic tools and workflows was implemented as the Metavisitor package in the Galaxy framework. Using the graphical Galaxy workflow editor, users with minimal computational skills can use existing Metavisitor workflows or adapt them to suit specific needs by adding or modifying analysis modules. Metavisitor can be used on our Mississippi server, or can be installed on any Galaxy server instance and a pre-configured Metavisitor server image is provided. Metavisitor works with DNA, RNA or small RNA sequencing data over a range of read lengths and can use a combination of de novo and guided approaches to assemble genomes from sequencing reads. We show that the software has the potential for quick diagnosis as well as discovery of viruses from a vast array of organisms. Importantly, we provide here executable Metavisitor use cases, which increase the accessibility and transparency of the software, ultimately enabling biologists or clinicians to focus on biological or medical questions.

## Introduction

Viruses infect cells and manipulate the host machinery for their replication and transmission. Genomes of viruses show high diversity and can consist of single- or double-stranded RNA or DNA. Many types of viral replication cycles exist which may involve various cellular compartments, various DNA or RNA replication intermediates,and diverse strategies for viral RNA transcription and viral protein translation. Next nbsp; generation deep-sequencing has become a powerful approach for virologists in their quest to detect and identify viruses in biological samples, even when they are present at low levels. However, none of the existing sequencing methods allows comprehensive detection of all virus classes. For instance, DNA sequencing fails to detect viruses lacking a DNA stage whereas commonly used mRNA sequencing protocols based on poly-A tailed RNA purification fail to detect viruses without poly-A tails.

Plants and invertebrates use RNA interference as an antiviral mechanism [1, 2]. Active antiviral RNAi results in significant enrichment of viral interfering small RNAs (viRNAs) relative to endogenous small RNAs (endo-siRNA). The ratio of viRNA reads over endo-siRNA reads depends on several factors such as the ability of a virus to replicate in the host and to evade the host RNAi machinery. Moreover, viRNAs derived from a variety of viruses can be detected in host organisms, regardless if these viruses have positive single strand, negative single strand or double-stranded RNA genomes,or DNA genomes [3]. Together, these features make small RNA deep sequencing a potent approach to detect viruses regardless of their genomic specificities, and different bioinformatic tools have been developed for detection or de novo assembly of viral genomes.

Accordingly, viRNAs produced by the insect model *Drosophila melanogaster* in response to viral infections were sufficient to reconstruct and improve the genomic consensus sequence of the Nora virus [4] using the Paparazzi perl script [5] which wraps the SSAKE assembler [6]. In this study, Paparazzi improved the consensus sequence and the coverage of the Nora virus genome by ∼20%, as compared to the previous Nora virus reference genome. SearchSmallRNA, a standalone tool with a graphical interface written in JAVA language, used a similar approach to reconstruct viral genomes [7]. Source 2 codes of both Paparazzi and SearchSmallRNA require specific skills for installation and execution as well as the retrieval of viral reference sequences. Furthermore neither program is currently available for download. Since both programs require known, closely related viral references for proper guidance of genome reconstructions from viRNAs, identification of more distant viral species or discovery of novel or unexpected viruses is precluded.

To circumvent the need for viral reference sequences, Velvet [8]de novo assemble contigs from plant [9], fruit fly and mosquito [10]have then been aligned to NCBI sequence databases, allowing the identification of partial or complete viral genomes. Several studies improved the strategy by combining two *de novo* assemblers [11–14], or scaffolding the contig pieces that could be blast-aligned to NCBI sequences using an additional translation-guided assembly step [15].

Collectively, the reported work allowed important progress in virus assembly and identification from deep sequencing data. However, the existing computational workflows are poorly accessible to a broad user base of biologists because they require specialist skills for installation, execution and adaptation to specific research. These skills may not be sufficient in some cases where tools are no longer available or documentation is missing.

In this context, we developed Metavisitor as a free and open source set of Galaxy tools and workflows [16, 17] allowing both *de novo* reconstruction of novel viruses and detection of already identified viral species from sequencing datasets. Using the graphical Galaxy workflow editor, Metavisitor workflows can be adapted to suit specific needs, by adding analysis steps or replacing/modifying existing ones. For instance, Metavisitor may help in field surveillance of insect vectors and emerging viral species during epidemics, in viral metagenomic studies or in experimental research or diagnosis for human patients suffering from viral infections or coinfections. In order to improve as much as possible the accessibility and usability of Metavisitor, we detailed a series of use cases that can be directly examined, replayed, tested or adapted using our Galaxy server (http://mississippi.fr). To ensure the sustainability of these executable use cases, a Galaxy server instance provisioned with the Metavisitor tools and workflows is also available as a Docker image. We expect that these tools will provide biologists and 3medical practitioners with an easy-to-use and adaptable software for the detection or identification of viruses from NGS datasets.

## Methods

Metavisitor consists of a set of Galaxy tools (Figure 1) that can be combined to (i) extract sequencing reads that do not align to the host genomes, known symbionts or parasites; (ii) retrieve up to date nucleotide as well as protein sequences of viral genomes deposited in Genbank [18] and index these sequences for subsequent blast, bowtie, etc. alignments; (iii) perform *de novo* assembly of extracted sequencing reads using Oases or Trinity, align the *de novo* contigs against the viral nucleotide or protein blast databases using blastn or blastx, respectively, and generate reports from blast outputs to help in known viruses diagnosis or in candidate virus discovery; (iv) use CAP3 (optional, see Use Case 3-3), blast and viral scaffolds for selected viruses to generate guided final viral sequence assemblies of blast sequence hits. For clarity, we group analysis steps below in functional tasks (i to iv). However, as shown in the Use Cases section, Metavisitor links these tasks in full workflows that can be executed once to generate complete and adapted analyses.

**Figure 1.**
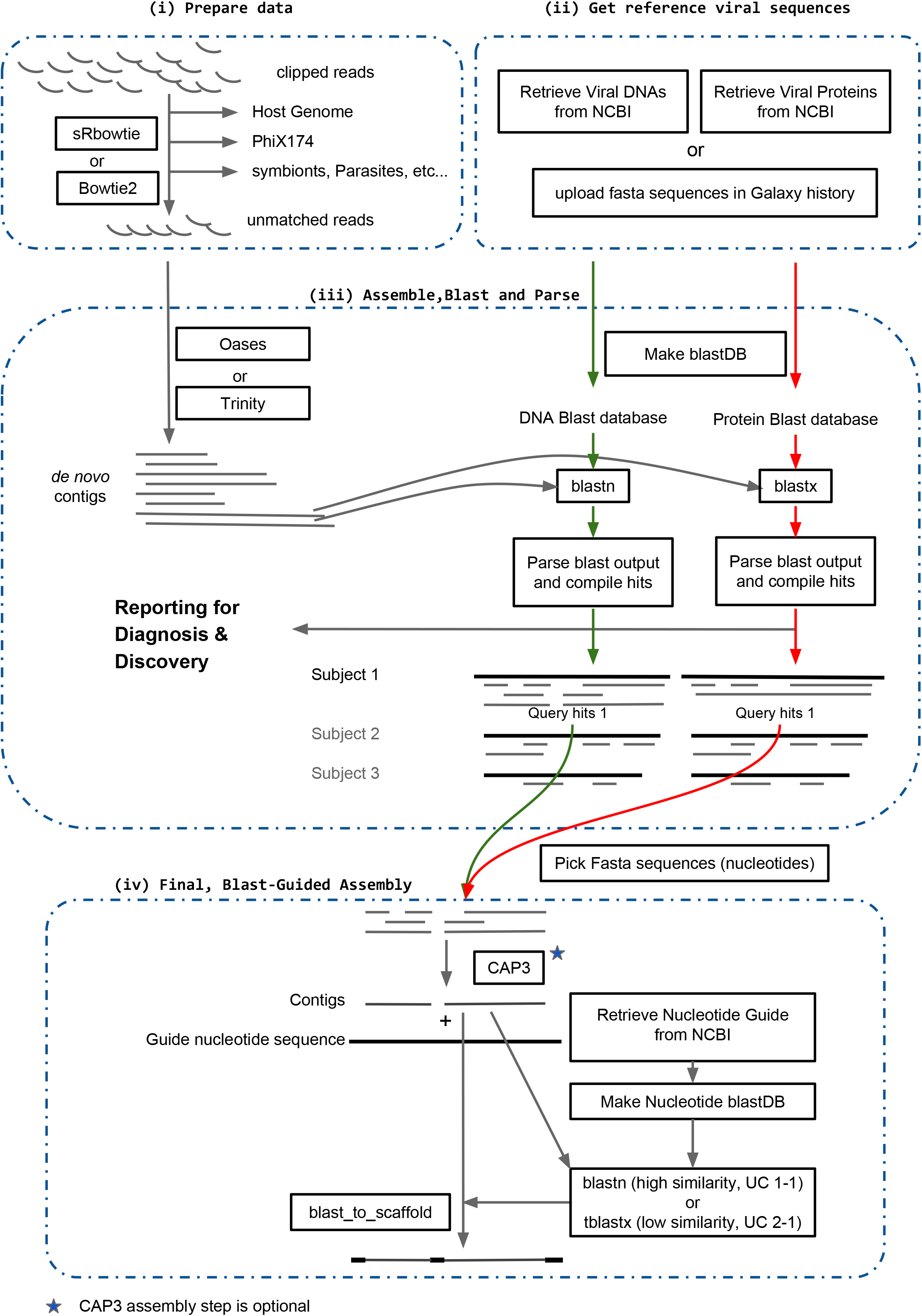
Global view of the Metavisitor workflow. The workflow is organised in sub workflows (dashed line) corresponding to functional tasks as described in the manuscript. All Galaxy Tools (square boxes) are available in the main Galaxy tool shed (https://toolshed.g2.bx.psu.edu/).

### (i) Prepare data

The purpose of the “Prepare data” task (Figure 1) is to process Illumina sequencing datasets in order to optimize the subsequent *de novo* assembly of viral sequencing reads. Raw sequence files in fastq or fasta format are clipped from library adapters and are converted to a weighted fasta file in which sequences are headed by a character string that contains a unique identifier and the number of times that the sequences were found in the dataset. These steps (which are optional) remove sequence duplicates and drastically reduces the workload of the next steps as well as the coverage variations after *de novo* assembly (see Use Cases 1-1 to 1-3). Datasets are then depleted from non-viral sequences by sequential alignments to the host genome, to other genomes from known or potential symbionts and parasites, as well as to PhiX174 genome sequences which are commonly used as internal controls in Illumina sequencing and may contaminate the datasets. The sequence reads that did not match the reference genomes are retained and returned.

### (ii) Get reference viral sequences

The “Get reference viral sequences” task is performed using the “Retrieve FASTA from NCBI” tool that sends a query string to the Genbank database [18] and retrieves the corresponding nucleotide or protein sequences. With this tool, a typical query for virus sequences retrieval is “txid10239[Organism] NOT txid131567[Organism] NOT phage”, which retrieves viruses sequences (txid10239) while filtering out cellular organisms sequences (txid131567) and phage sequences. This query was submitted to the nucleotide or protein Genbank databases (oct 2015) to retrieve the viral nucleotide and protein databases referred to as “vir1” in the rest of the article. However, users can change the tool settings by entering query strings that fit their specific needs. As downloading large sequence datasets from NCBI may take several hours with this query, we allow users to skip it by directly accessing the nucleotides or protein vir1 datasets on the Mississippi server (http://mississippi.fr) or to download it from figshare (https://dx.doi.org/10.6084/m9.figshare.3179026). For convenience, nucleotide and protein blast indexes are also available in the public library of the Mississippi server (but can also be generated using the “NCBI BLAST+ makeblastdb” Galaxy tool). Bowtie as well as bowtie2 indexes of the vir1 nucleotide sequences have been generated in the Mississippi Galaxy instance using the corresponding “data manager” Galaxyt ools. Finally, users can upload their own viral nucleotide and protein sequences using ftp and transfer them in a Galaxy history (Figure1).

### (iii) Assemble, Blast and Parse

In the task “Assemble, Blast and Parse”, RNA sequences returned by the “Prepare data” task are subjected to several rounds of *de novo* assembly by Velvet [8] using the Oases software package [19] and k-mer lengths ranging from 15 to 35 (for small RNA sequences) or from 13 to 69 (for longer RNA sequences). Importantly, as illustrated in Use Case 3-3 (see below), the Oases assembler can be replaced by a different *de novo* assembler such as Trinity [20] that performs better with longer sequencing reads.

In a next step, *de novo* assembled contigs are aligned to both “virus nucleotides” and “virus proteins” vir1 BLAST databases built from the viral reference sequences (Figure 1) using the blastn or blastx Galaxy tools [21] that search nucleotide or protein databases using nucleotide or translated nucleotide queries, respectively [22]. Default parameters for these tools are adjusted in order to report only the 5 best alignments per contig (Maximum hits set to 5) and to generate a tabular blast output that includes the 12 standard columns plus a column containing the length of the aligned subject sequences (extended columns, “slen” checked).

Tabular outputs generated by blastn and blastx alignments are next parsed by the “Parse blast output and compile hits” tool to return 4 files. In the “blast analysis, by subjects” output file (Supplementary Figure 1), the subject sequences in the viral nucleotide or protein blast databases that produced significant blast alignments (hits) with Oases contigs are listed, together with those contigs and the blast information associated to the hits (% Identity, Alignment Length, start and end coordinates of hits relatively to the subject sequence, percentage of the contig length covered by the hit, E-value and Bit Score of the hit). In addition, for each subject sequence in the list, the length in nucleotide or amino-acid of the subject sequence (Subject Length), the summed coverage of the subject by all contig hits (Total Subject Coverage) as well as the fraction of the subject length that this coverage represents (Relative Subject Coverage), and the best (Best Bit Score) and mean (Mean Bit Score) bit scores produced by contig hits are computed and indicated. A simplified output can be generated without contigs and blast information by using the “compact” option for the reporting mode of the “Parse blast output and compile hits” tool. A second “hits” output file generated by the tool contains the sequences of contig portions that produced significant alignment in the BLAST step (i.e. query hit sequences), flanked by additional contig nucleotides 5’ and 3’ to the hit (the size of these margins is set to 5 by default and can be modified by the user). Finally, the tool returns the contigs that produced significant blast hits (“Blast aligned sequences”) as well as those that did not (“Blast unaligned sequences”).

### (iv) Final assembly from blastn or blast

The last “Final assembly from blastn/x” task (Figure 1) allows to manually choose candidates from user’s inspection of the “blast analysis, by subjects” file and to generate further sequence assembly. Using the tool “Pick Fasta sequences” and the appropriate query string, users first retrieve from the file “hits” all blastn or blastx hits that significantly matched a subject sequence. When necessary, these hit sequences can be further assembled in longer contigs using the “cap3 Sequence Assembly” Galaxy tool adapted from CAP3 [23]. In some cases (see below), unique viral contigs can already be obtained at this step. In cases where there are still multiple unlinked contigs, the workflow provides the possibility to generate a single composite sequence where these contig sequences (indicated in uppercase characters) are integrated in a matched subject sequence taken as a scaffold (lowercase characters). This is done by (a) retrieving the subject sequence from the NCBI nucleotide databases, generating a blast nucleotide index from this sequence and aligning the contigs to this index with blastn or tblastx tools, and (b) running the “blast_to_scaffold” tools by taking as inputs the contigs, the guide/scaffold sequence and the blastn or blastx output (Figure 1, bottom).

## Availability of Metavisitor

All Metavisitor tools and workflows are installed in the Galaxy server http://mississippi.snv.jussieu.fr. Readers can easily review all use cases described below by following the indicated html links to this server. Moreover, they can also import in their personal account the published Metavisitor use case histories and their corresponding workflows to re-run the described analyses or adapt them to their studies.

We made all tools and workflows that compose Metavisitor available from the main Galaxy tool shed (https://toolshed.g2.bx.psu.edu/), in the form of a tool suite (suite_metavisitor_1_2) which thus can be installed and used on any Galaxy server instance. The Metavisitor workflows are also available from the *myexperiment* repository (http://www.myexperiment.org/) They can be freely modified or complemented with additional analysis steps with in the Galaxy environment.

The Metavisitor tool codes are accessible in our public GitHub repository (https://github.com/ARTbio/tools-artbio/). We also provide a Docker image artbio/metavisitor:1.2 as well as an ansible playbook that both allow to deploy a Galaxy server instance with preinstalled Metavisitor tools and workflows in local infrastructures. Extensive documentation on how to install and use Metavisitor Galaxy servers is available at https://artbio.github.io/GalaxyKickStart/about_metavisitor.

## Results/Use Cases

In this section, we present use cases to demonstrate the use of various Metavisitor workflows adapted to specific situations and dataset formats. For each use case, we briefly present the purpose of the original study from which the datasets originate, and we propose html links to input data, workflows as well as to histories generated with these workflows and input data. Using this process, we intend to provide transparent and executable analyses: readers can examine the use cases in every detail using the Galaxy web interface; they can also import the input data, histories and workflows in their own Galaxy Mississippi account and re-execute the analyses as we did; finally they can experiment Meta visitor work flows with their own datasets and parameters.

### 1. Detection of known viruses

#### Use Cases 1-1, 1-2 and 1-3

Using small RNA sequencing libraries SRP013822 (EBI ENA) and the Paparazzi software [5] we were previously able to propose a novel reference genome (NCBI JX220408) for the Nora virus strain infecting Drosophila melanogaster stocks in laboratories [4]. This so-called rNora genome differed by 3.2% nucleotides from the Nora virus reference NC_007919.3 and improved the alignment rate of viral siRNAs by ∼121%. Thus, we first tested Metavisitor on the small RNA sequencing datasets SRP013822.

Three Metavisitor workflows were run on the merged SRP013822 small RNA sequence reads and the NC_007919.3 genome as a guide for final reconstruction (in Galaxy history “Input data for Use Cases 1-1, 1-2, 1-3 and 1-4”). The first Workflow for Use Case 1-1 used raw reads collapsed to unique sequences (see methods section) to reconstruct a Nora virus genome referred to as Nora_MV (dataset 35) in the History for Use Case 1-1. In a second Workflow for Use Case 1-2, we did not collapse the SRP013822 reads to unique sequences (see materials and method), which allowed the reconstruction of a Nora_raw_reads genome (dataset 33) in the History for Use Case 1-2. In a third Workflow for Use Case 1-3, the abundances of SRP013822 sequence reads were normalized using the Galaxy tool “Normalize by median” [24], which allowed the reconstruction of a Nora_Median-Norm-reads genome (dataset 37) in the History for Use Case 1-3.

All three reconstructed genomes as well as the Paparazzi-reconstructed JX220408 genome had a high sequence similarity (>96.6% nucleotide identity) with the NC_007919.3 guide genome (see Supplementary File 1). The final *de novo* (capital letters) assemblies of both the Nora_raw_reads and Nora_Median-Norm-reads genomes entirely covered the JX220408 and NC_007919.3 genomes (both 12333 nt), whereas the *de novo* assembled part of the Nora_MV genome was marginally shorter (12298nt, the 31 first 5’ nucleotides are in lowercase to indicate that they were not *de novo* assembled but instead recovered from the guide genome). To evaluate the quality of assemblies, we use the “Workflow for remapping in Use Cases 1-1,2,3” (from the history Input data for Use Cases 1-1, 1-2, 1-3 and 1-4) for remapping of the SRP013822 raw reads to the 3 reconstituted Nora virus genomes as well as to the JX220408 guide genome. As can be seen in “History for remapping in Use Cases 1-1,2,3” and Fig. 2, SRP013822 reads matched the genomes with almost identical profiles and had characteristic size distributions of viral siRNAs with a major peak at 21 nucleotides. Importantly, the numbers of reads re-matched to the Nora virus genomes were 1,578,704 (Nora_MV) > 1,578,135 (Paparazzi - JX220408) > 1,566,909 (Nora_raw_reads) > 1,558,000 (Nora_Median-Norm-reads) > 872,128 (NC_007919.3 reference genome guide).

**Figure 2.**
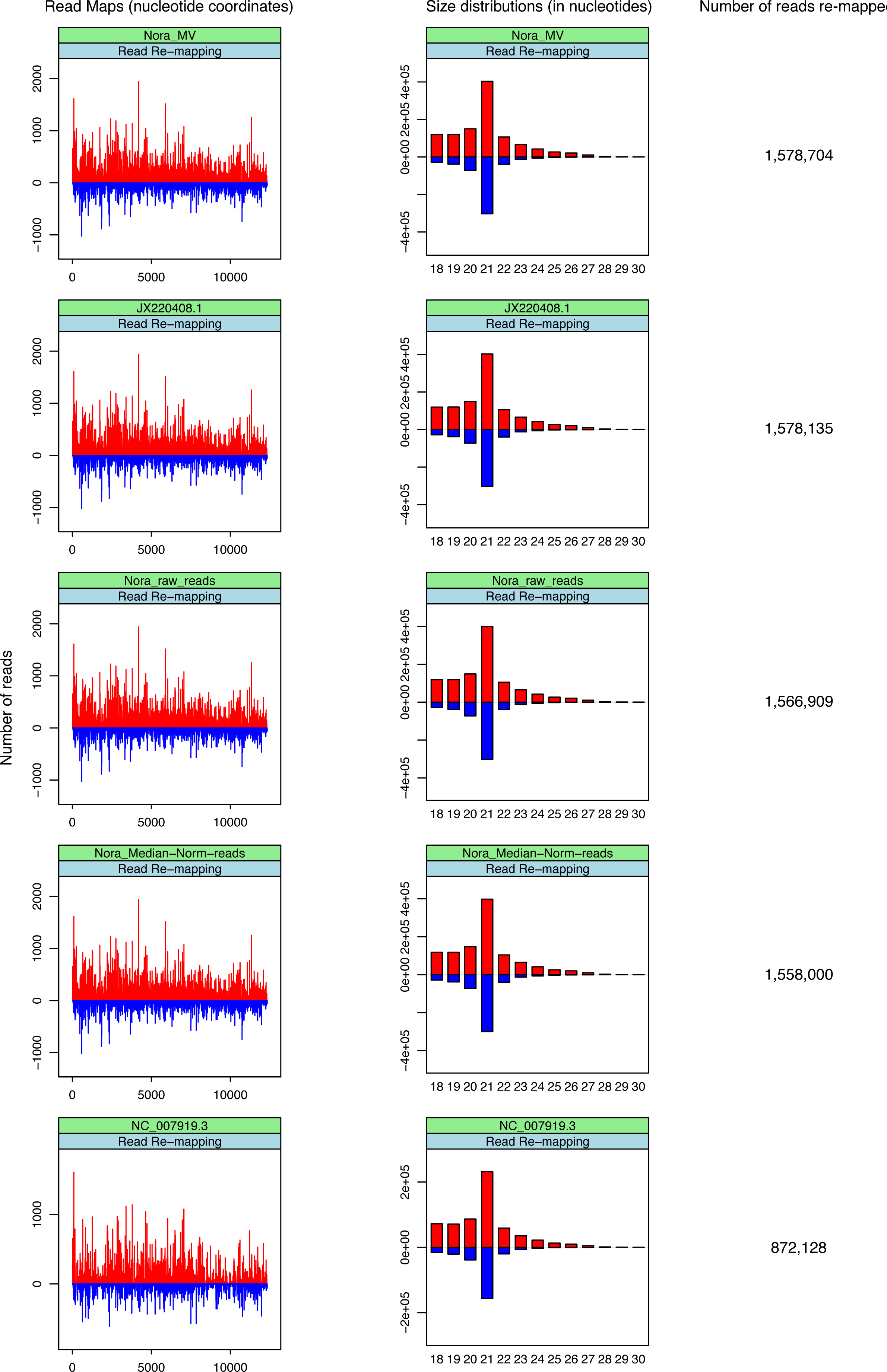
Realignments of small RNA sequence reads to reconstructed (Nora_MV, Nora_raw_reads and Nora_Median−Norm−reads) or published (JX220408.1 and NC_007919.3) Nora virus genomes. Plots (left) show the abundance of 18–30-nucleotide (nt) small RNA sequence reads matching the genome sequences and histograms (middle) show length distributions of these reads. Positive and negative values correspond to sense and antisense reads, respectively. Total read counts are indicated to the right hand side.

Thus, Metavisitor reconstructed a Nora virus genome Nora_MV whose sequence maximizes the number of vsiRNA read alignments which suggests it is the most accurate genome for the Nora virus present in the datasets. Of note, the Nora_MV genome differs from the JX220408 rNora genome generated by Paparazzi by only two mismatches at positions 367 and 10707, and four 2nt-deletions at positions 223, 365, 9059 and 12217 (see Supplementary File 1). These variations did not change the amino acid sequence of the 4 ORFs of the Nora virus. We conclude that Metavisitor performs slightly better than Paparazzi for a known virus, using *de novo* assembly of small RNA reads followed by blast-guided assembly. We did not observe any benefits of using raw reads (Metavisitor Use Case 1-2) or normalized-by-median reads (Metavisitor Use Case 1-3) for the Oases assembly, but rather a decrease in the accuracy of the reconstructed genome as measured by the number of reads re-mapped to the final genomes (Figure2).

#### Use Case 1-4

In order to show the ability of Metavisitor in detecting multiple known viruses in small RNA sequencing datasets, we derived a simplified workflow from the “Workflow for Use Case 1-1”, where blastn alignments of Oases contigs are simply parsed using the “Parse blast output and compile hits” tool without any filtering. Using this Workflow for Use Case 1-4 with the SRP013822 sequence datasets as input returned a list of parsed blastn alignments in the “History for Use Case 1-4” which contains, as expected, the Nora virus. In addition, Oases contigs were found to align with high significance (Mean BitScore > 500) to the Drosophila A virus and to the Drosophila C virus (Dataset 18 and Table 1), strongly suggesting that the fly stocks analyzed in our previous work were also subject to persistent infection by these viruses [4].

**Table 1.**
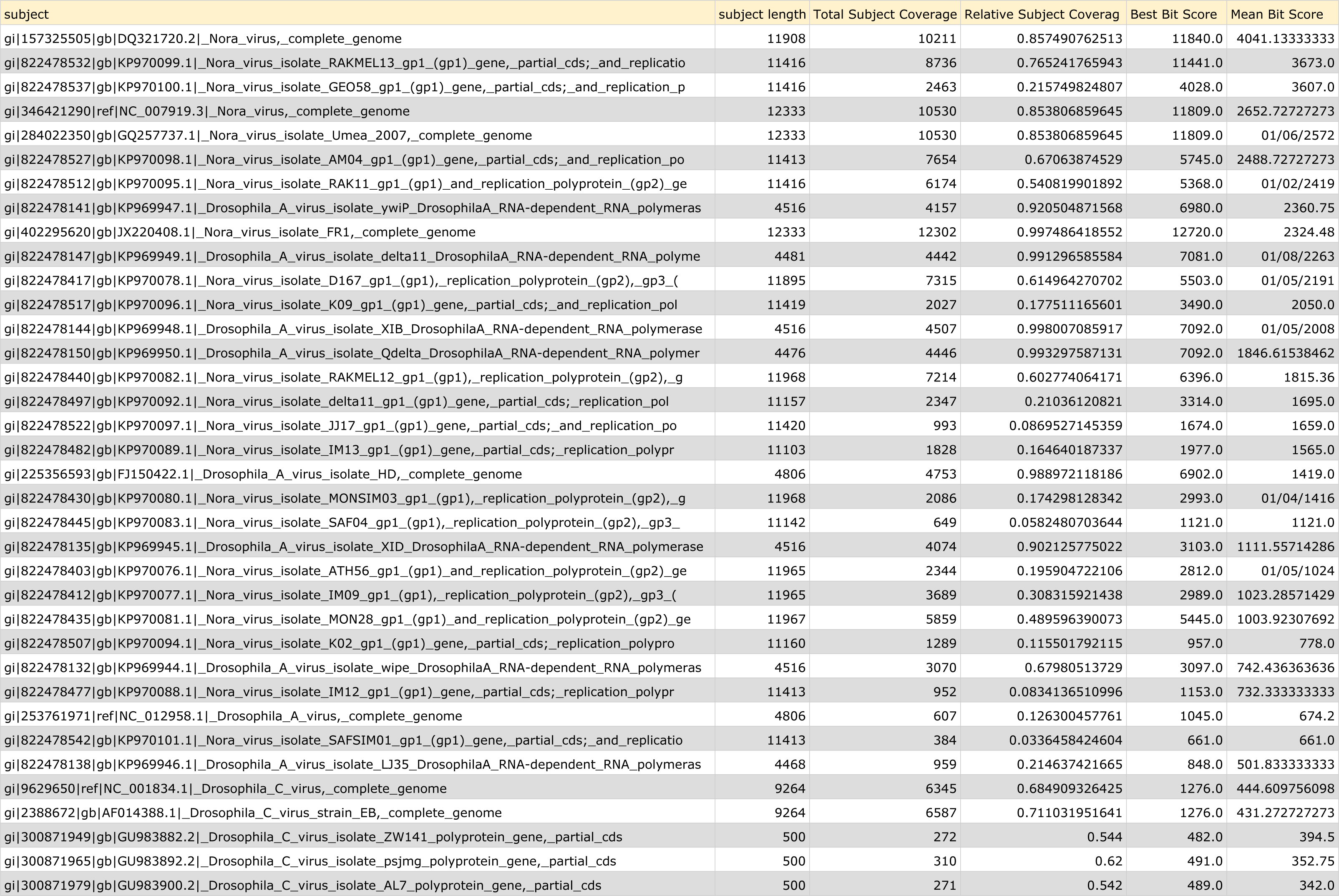
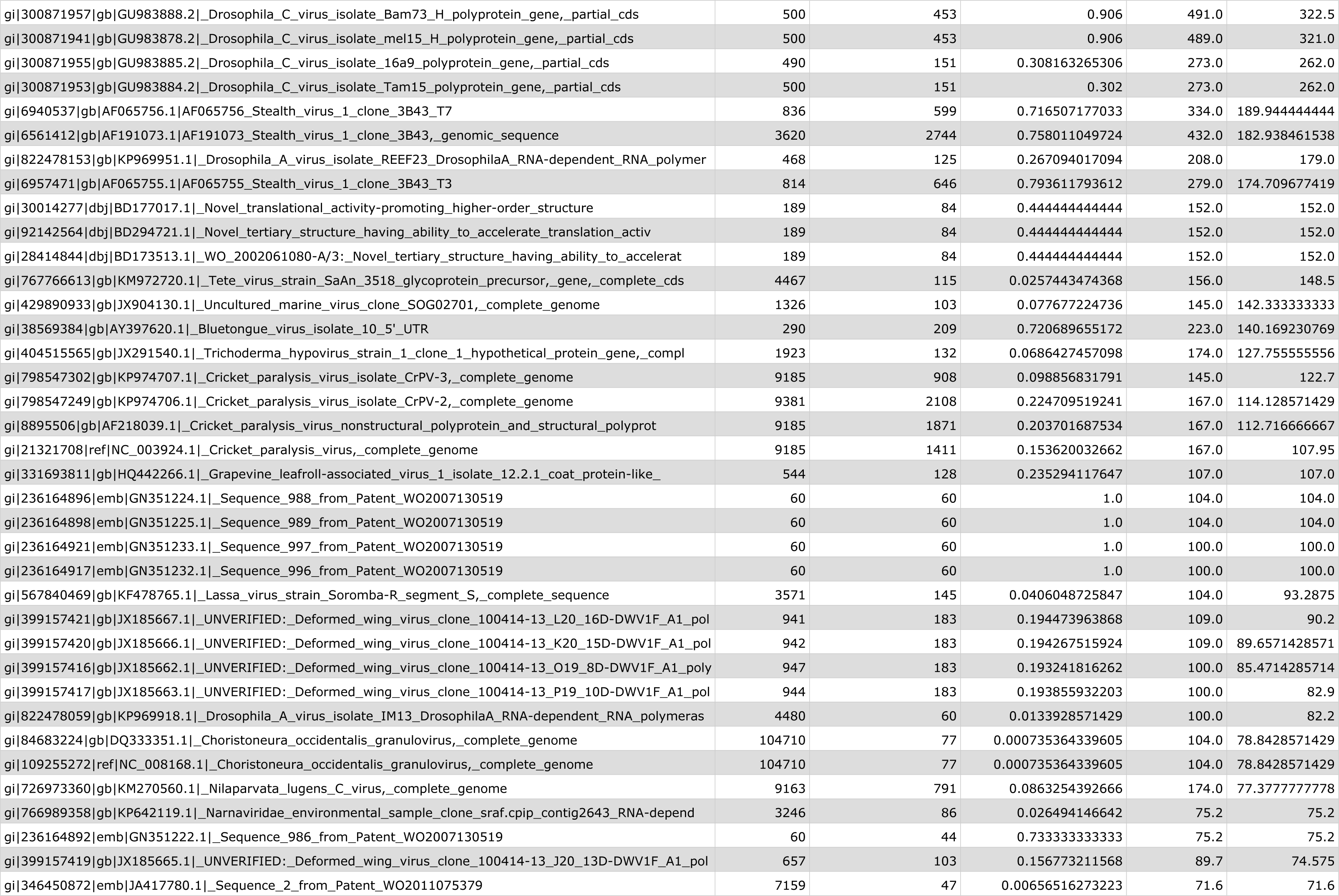
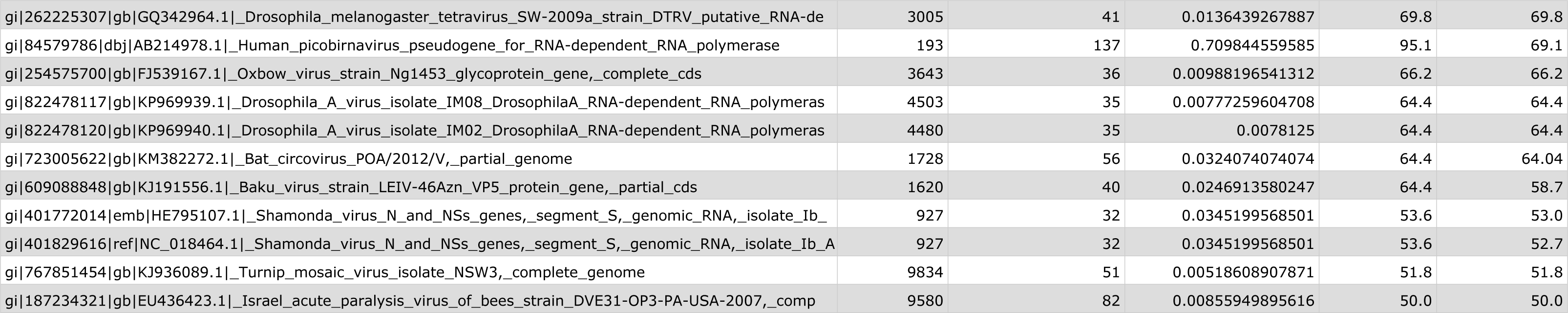
Report table generated by the “Parse blast output and compile hits” tool in History for Use Case 1-4 showing the presence of *Drosophila* A virus and *Drosophila* C virus in addition to the Nora virus in the small RNA sequencing of laboratory *Drosophila*. See Method section for a description of the columns.

### 2. Discovery of novel viruses

#### Use Case 2-1

We recently discovered two novel viruses infecting a laboratory colony of *Anopheles coluzzii* mosquitoes [25]. Using small RNA datasets from these mosquitoes (study accession number ERP012577, history “Input data for Use Cases 2-1 and 2-2”), and the Workflow for Use Case 2-1, we were able to assemble a number a Oases contigs that showed significant blastx hits with *Dicistroviridae* proteins, including *Drosophila* C virus (DCV) and Cricket paralysis virus (CrPV) proteins (see the dataset 26 produced by the “Parse blast output and compile hits” tools in the History for Use Case 2-1. The viral family of *Dicistroviridae* was named from the dicistronic organisation of their genome. A 5’ open reading frame codes for a non-structural polyprotein and a second non-overlapping 3’ open reading frame codes for the structural polyprotein of the viral particle.

In order to construct a potential new *A. coluzzii* dicistrovirus genome, we thus collected blastx hits showing significant alignment with both *Drosophila* C virus and *Cricket* paralysis viral polyproteins (dataset 32, Dicistroviridae Hits), and we further assembled these hits using CAP3, which produced 4 contigs of 1952, 341, 4688 and 320 nt,respectively. We then aligned these 4 contigs to the DCV genome NC_001834.1 with tblastx and used the “blast_to_scaffold” tool to produce a final assembly (dataset 42: “New AnCV sequences in DCV scaffold”). Re-mapping of the ERP012577 small RNA reads using the Workflow for remapping in Use Cases 1-1,2,3 adapted to ERP012577 at runtime showed that they mostly align to *de novo* assembled regions (uppercase nucleotides) of this chimeric genome and have a typical size distribution of viral derived siRNA (see dataset 64), suggesting that the NC_001834.1 DCV sequences of the scaffold (lowercase nucleotides) are loosely related to the actual sequence of the novel *Anopheles coluzzii* dicistrovirus. Nevertheless, the composite assembly already allows designing primers in the *de novo* assembled regions to PCR amplify and sequence the regions of the viral genome that could not be *de novo* assembled.

#### Use Case 2-2

We next used RNAseq libraries from the same *Anopheles coluzzii* colony available in the history Input data for Use Cases 2-1 and 2-2 (dataset 19, deposited in EBI-SRA under accession number ERS977505) to demonstrate the use of a Metavisitor workflow with long RNA sequencing read datasets. Thus, to generate the Galaxy History for Use Case 2-2 with the Workflow for Use Case 2-2, 100nt reads were aligned without any clipping to the *Anopheles gambiae* genome using bowtie2, and unmatched read were subjected to Oases assembly (kmer range, 25 to 69). Oases contigs were then filtered for a size > 5000 nt and aligned to the protein viral reference using blastx. Parsing of blastx alignments with the “blast analysis, by subjects” tool repeatedly pointed to a 8919nt long Oases contig that matched to structural and non-structural polyproteins of DCV and CrPV (dataset 24 in history for Use Case 2-2). This 8919nt contig (dataset 29 in History for Use Case 2-2) completely includes the contigs generated with the small RNA datasets and shows a dicistronic organization which is typical of Dicistroviridae and is referred to as a novel *Anopheles* C Virus [25]. The sequence of this *Anopheles* C Virus is deposited to the NCBI nucleotide database under accession number KU169878. As expected, the ERP012577 small RNA reads realigned to this genome (using the Workflow for remapping in Use Cases 2-1,2) now show a typical alignment profile all along the AnCV genome sequence with a size distribution peaking at the 21nt length of viral derived siRNAs and no gap (dataset 84: Size distribution and Readmaps in Galaxy history Metavisitor Use Case2-2).

Taken together, the Metavisitor Use Cases 2-1 and 2-2 illustrate that when short read datasets do not provide enough sequencing information, a simple, adapted Metavisitor workflow is able to exploit long reads of RNA sequencing datasets, if available, to assemble a complete viral genome.

### 3. Virus detection in human RNA seq libraries

Having illustrated that Metavisitor is able to generate robust genome assemblies from known and novel viruses in *Drosophila* and *Anopheles* sequencing datasets, we tested whether it can be used as a diagnostic workflow to detect viruses in RNA sequencing datasets of human patients from three different studies [26–28].

#### Use Case 3-1

Innate lymphoid cells (ILCs) play a central role in response to viral infection by secreting cytokines crucial for immune regulation, tissue homeostasis, and repair. Therefore, the pathogenic effect of HIV on these cells was recently analyzed in infected or uninfected patients using various approaches, including transcriptome profiling [27]. ILCs are unlikely to be infected *in vivo* by HIV as they lack expression of the CD4 co-receptor of HIV and they are refractory *in vitro* to HIV infection. However, we reasoned that ILCs samples could still be contaminated by infected cells. This might allow Metavisitor to detect and assemble HIV genomes from patient’s ILC sequencing data (EBI SRP068722). As these datasets contains short 32 nt reads which in addition had to be 3’ trimmed to 27 nt to retain acceptable sequence quality, we designed a Workflow for Use Case 3-1 that is similar to the workflows used in cases 1-1 and 2-1 for small RNA sequencing data. In that workflow however, sequencing datasets are depleted from reads aligning to the human genome (hg19) and viral reads are selected by alignment to the NCBI viral sequences using our sRbowtie tool. These reads are further submitted to Oases assembly (kmers 11 to 27), the resulting contigs are aligned to the Nucleotide Viral Blast Database using blastn. Alignments are parsed using the “Parse blast output and compile hits” tool, removing alignments to NCBI sequences related to patents to simplify the report (“Patent” term in the filter option of the “Parse blast output and compile hits” tool). Finally, a report is generated by concatenating the reports produced by this tool for each patient.

Using the Galaxy tool “Extract reads in FASTQ/A format from NCBI SRA”, we imported 40 sequence datasets from the EBI SRP068722 archive in the history “Input data for Use Case 3-1” and we merged the ICL datasets belonging to the same patients (datasets 43 to 59). We then generated a dataset collection of these patient sequence data (Patient collection) and executed the Workflow for Use Case 3-1 to perform all-in-one batch analysis of this collection in the History for Use Case 3-1 (summarized in Table 2). In this history analysis, we were able to detect HIV RNAs in samples from 3 out of 4 infected patients whereas all samples from control uninfected patients remained negative for HIV. This Metavisitor workflow was able to accurately detect HIV RNA, even in samples where the number of sequence reads was expected to be low,as mentioned above.

**Table 2.**
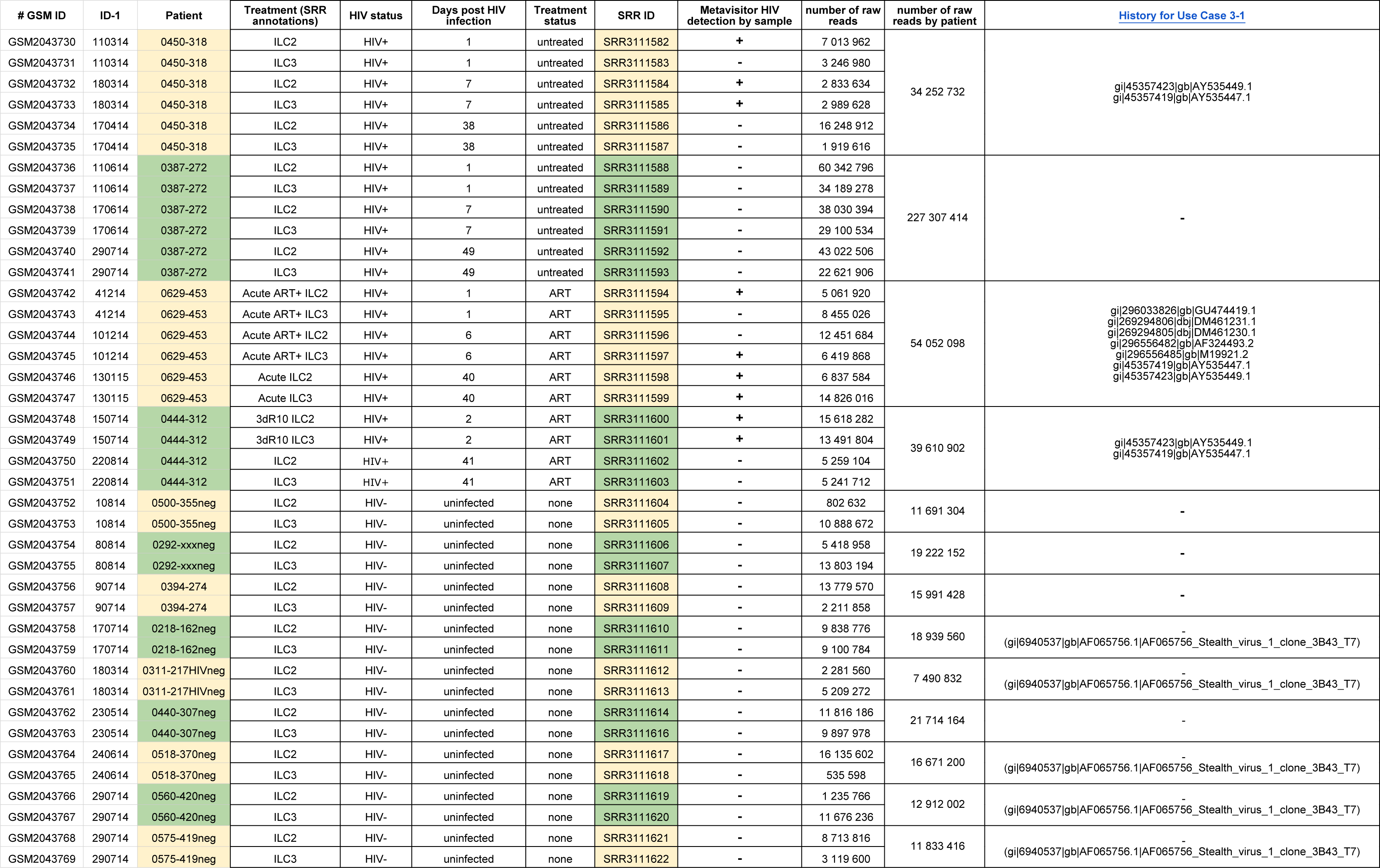
HIV detection in RNA sequencing datasets from ILC patient samples. The table summarizes the report generated by Metavisitor from a batch of 40 sequence datasets using the Workflow for Use Case 3-1 in the Galaxy History for Use Case 3-1(dataset 199). This table reports the metadata associated with each indicated sequence dataset as well as the ability of Metavisitor to detect HIV in datasets and patients.

#### Use Case 3-2

Yozwiak *et al.* searched the presence of viruses in RNA Illumina sequencing data from serums of children suffering from fevers of unknown origins [26]. In this study, paired-end sequencing datasets were depleted from reads aligning to the human genome and the human transcriptome using BLAT and BLASTn, respectively, and the remaining reads were aligned to the NCBI nucleotide database using BLASTn. A virus was considered identified when 10 reads or more aligned to a viral genome which was not tagged as a known lab contaminant.

For a significant number of Patient IDs reported in table 1 of the article [26], we were not able to find the corresponding sequencing files in the deposited EBI SRP011425 archive. In addition, we did not find the same read counts for these datasets as those indicated by the authors. With these limitations in mind, we used the Galaxy tool “Extract reads in FASTQ/A format from NCBI SRA” to download in the Galaxy history “Input Data for Use Case 3-2” 86 sequencing datasets that could be further concatenated and assigned to 36 patients in Yozwiak *et al* [26] (dataset collection 191). It is noteworthy that sequence reads in SRP011425 datasets are 97 nt long. Thus, the “Workflow for Use Case 3-2” that we built to perform all-in-one batch analysis is adapted from the Workflow for Use Case 3-1 with the following modifications: (i) sequences reads are depleted from human sequences and viral reads are selected by alignment to the NCBI viral sequences using the Galaxy bowtie2 tool instead of our sRbowtie tool; (ii) viral reads are submitted to Oases assembly using kmer values ranging from 13 to 69; (iii) the SAM file with reads alignments to the vir1 bowtie2 index is parsed using the “join” and “sort” Galaxy tools in order to detect putative false negative datasets with viral reads that fails to produce significant Oases viral contigs.

We executed the Workflow for Use Case 3-2 on the datasets from the history “Input Data for Use Case 3-2” to produce the History for Use Case 3-2. The information generated in this history is summarized in Table 3 (see also the dataset 484 “Virus identification by patient”) and shows that under these settings, Metavisitor detected the same viruses as those reported by Yozwiak *et al.* in 17 patients. Although viral reads were detected in 16 other patients, they were not covering sufficient portions of viral genomes to produce significant viral assemblies. Finally, in the three remaining patients (patients 363, 330 and 345 in Table 3 and corresponding Galaxy datasets datasets 424, 384 and 368), we detected viruses (Dengue virus 2, Stealth virus 1 and Dengue virus 4, respectively) other than those identified by Yozwiak *et al.* As mentioned above, these discrepancies are most likely due to misannotation of deposited datasets, which precludes further detailed comparisons.

**Table 3.**
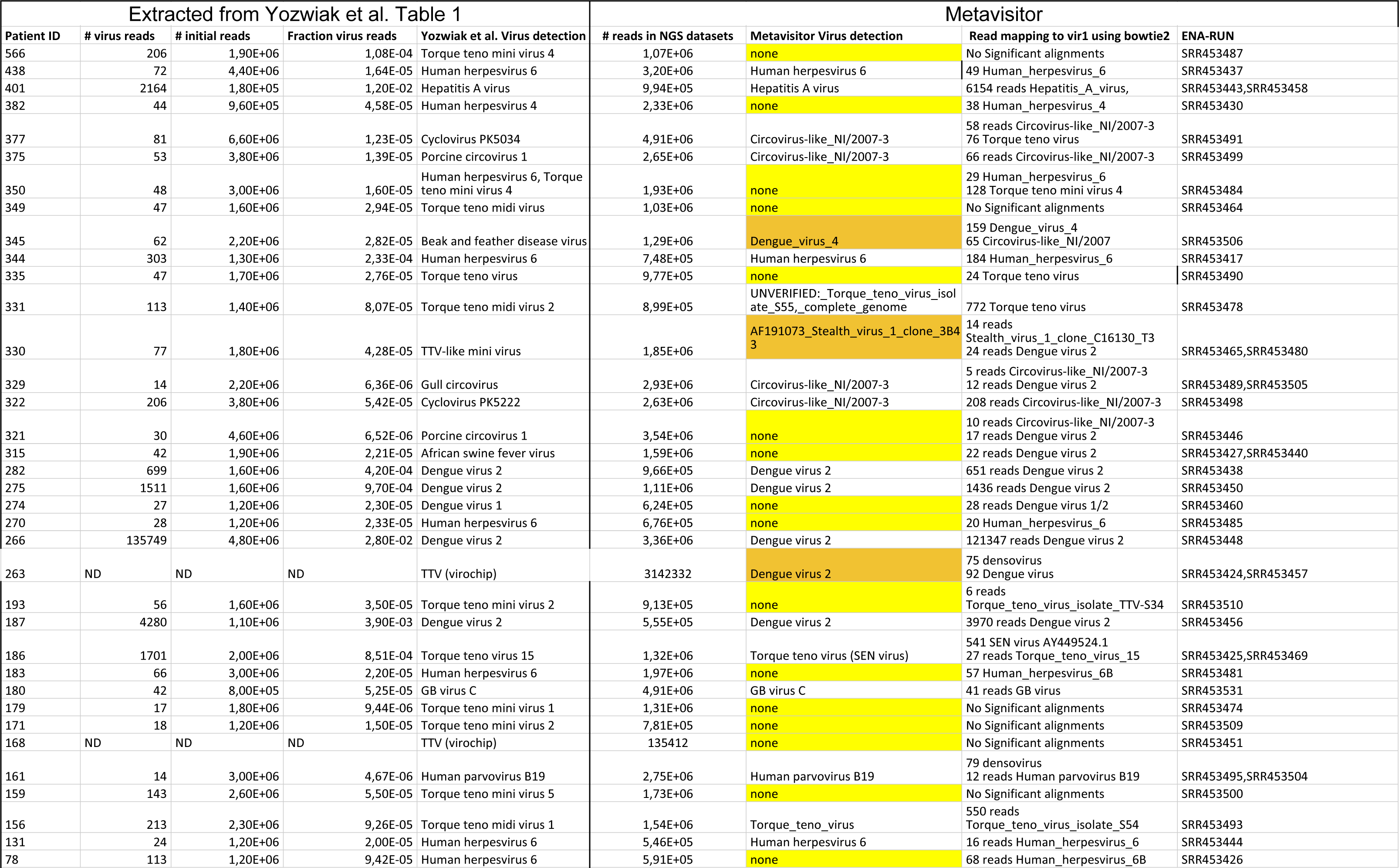
Virus detection in RNAseq datasets from 36 traceable patients by Metavisitor after blast alignment of viral contigs (dataset 484 in Galaxy History for Use Case 3-2). For detection of false positives, reads were aligned to the bowtie2 vir1 index before *de novo* assembly (see dataset collection 261 in the history). The data from this computational treatment are summarized in the column “Read mapping to vir1 using bowtie 2”and detailed ine ach corresponding patient tabs.

#### Use Case 3-3

Matranga *et al.* recently improved library preparation methods for deep sequencing of Lassa and Ebola viral RNAs in clinical and biological samples [28]. Accordingly, they were able to generate sequence datasets of 150 nt reads providing high coverage of the viral genomes. We used these datasets, relevant in the context of Lassa and Ebola outbreak and epidemic response, to demonstrate the versatility of Metavisitor as well as its ability to generate high throughput reconstruction of viral genomes.

In order to take into account the longer reads and higher viral sequencing depths in the available datasets [28], we adapted a Metavisitor workflow for Use Case 3-3 as follows: (i) The sequencing reads are directly aligned to the viral NCBI sequences without prior depletion by alignment to the human or rodent hosts; (ii) the Trinity *de novo* assembler[20] that performs well with longer reads is used instead of Oases; (iii) reconstruction of Lassa and Ebola genomes from the sequences of the blast hits with the nucleotide viral blast database is directly performed with our blast to scaffold tool without CAP3 assembly since the Trinity contigs are already covering a significant part of the viral genomes; (iv) the reports generated by our “Parse blast output and compile hits” tool as well as the reconstructed genome generated for each sample are concatenated in single datasets for easier browsing and subsequent phylogenetic or variant analyses; (v) finally, for adaptability to any type of virus, two input variables are specified by the user at the workflow runtime: the name of the virus to be searched for in the analysis, and the identifier of the sequenceto be usedas guide in genome reconstruction steps.

We imported 63 sequence datasets available in the EBI SRA PRJNA254017 and PRJNA257197 archives [28] in the history “Input Data for Use Case 3-3”, and grouped these datasets in Lassa virus (55 fastq files) and Ebola virus (8 fastq files) dataset collections (see Table 4 for the complete description of the analyzed samples). To generate the History for Use Case 3-3 Lassa L, we then executed the workflow for Use Case 3-3, taking the Lassa virus dataset collection as input sequences, “Lassa” as a filter term for the “Parse blast output and compile hits” tool and the NCBI sequence NC_004297.1 as a guide for reconstruction of the Lassa virus segment L. We also generated the History for Use Case 3-3 Ebola with the same workflow, taking the Ebola virus dataset collection as input sequences, “Ebola” as a filter term for the “Parse blast output and compile hits” tool and the NCBI sequence NC_002549.1 as a guide for reconstruction of the Ebola virus genome.

**Table 4.**
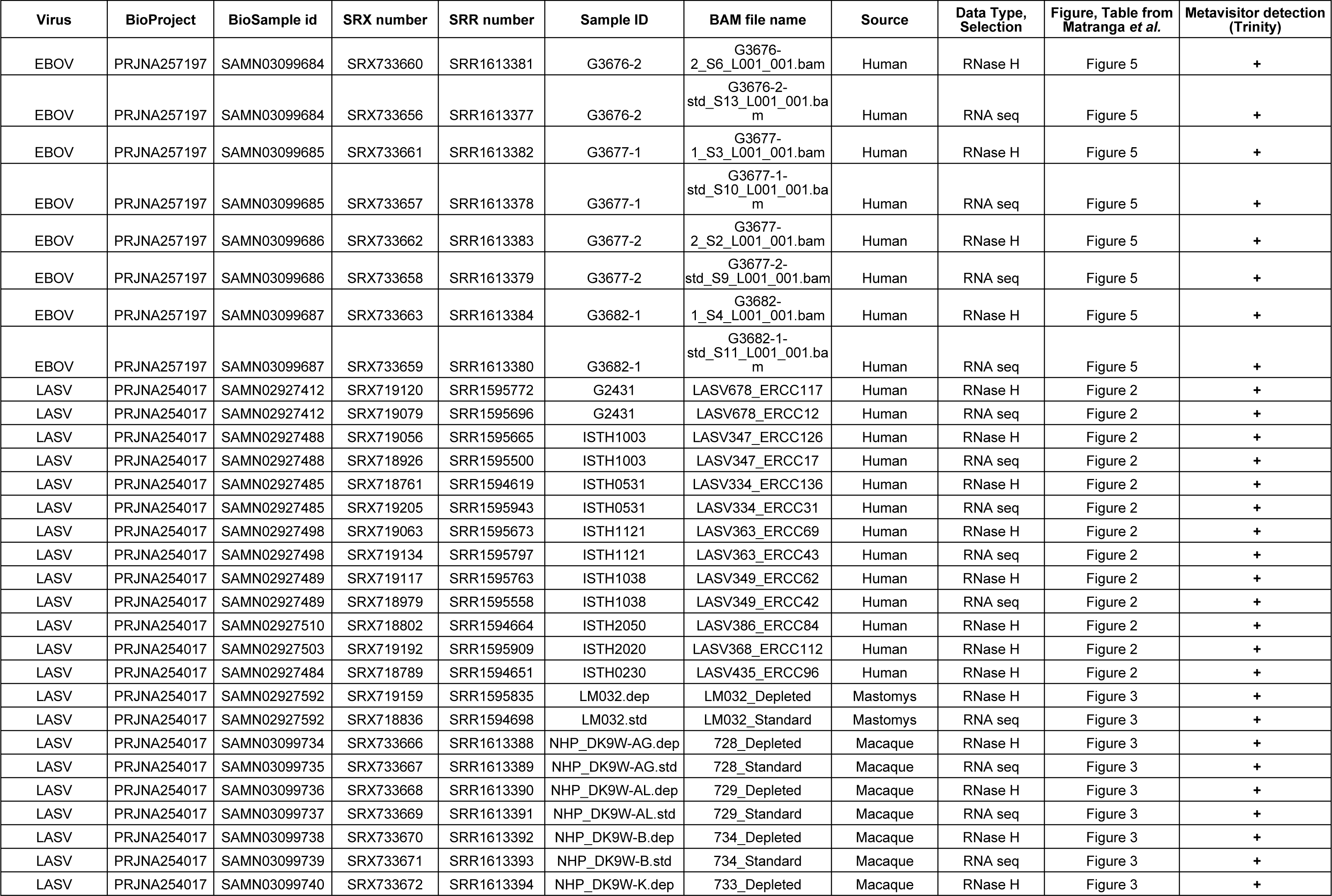
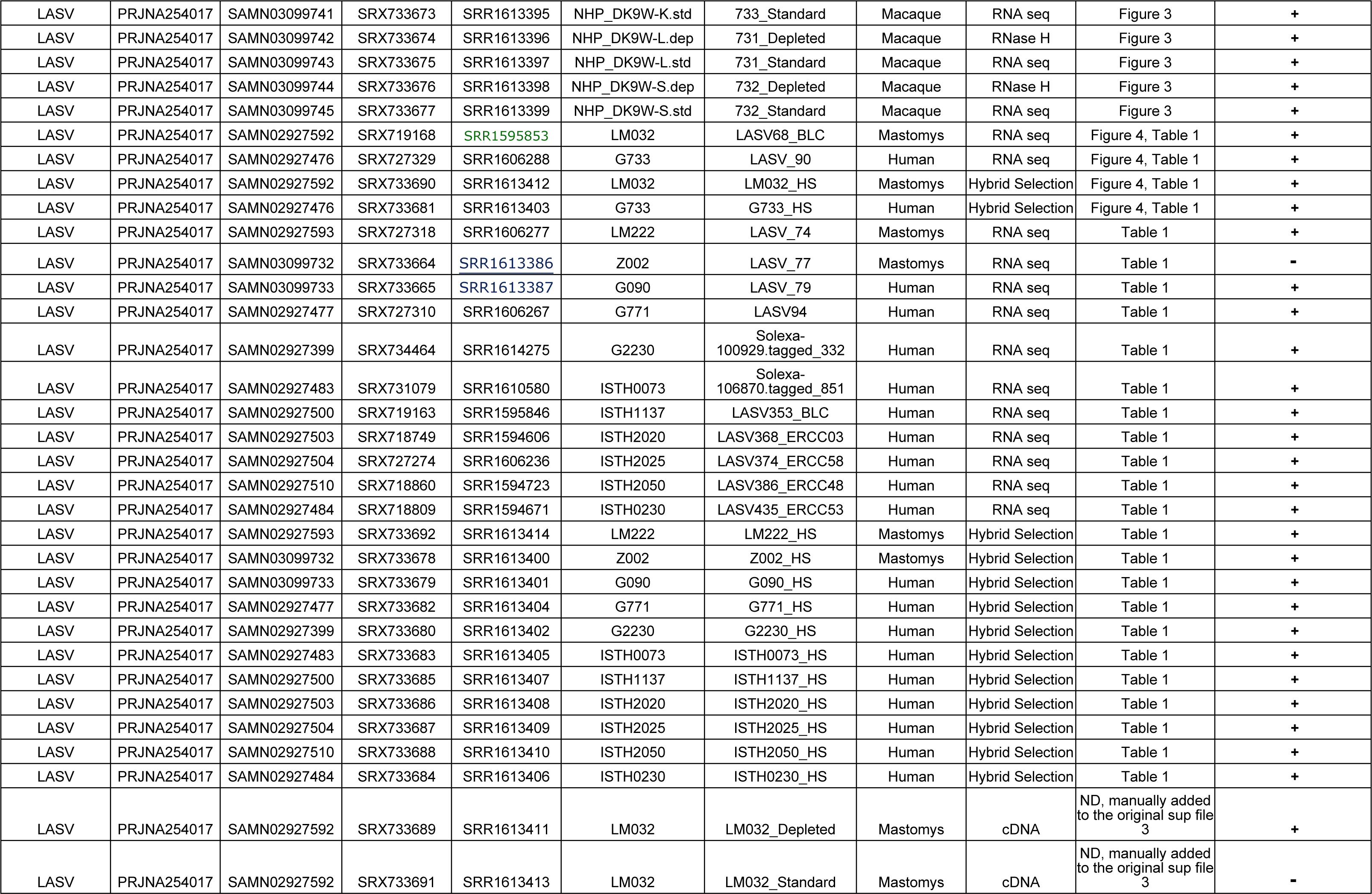
Metavisitor detection of Lassa virus (55 RNAseq datasets) and of Ebola virus (8 RNAseq datasets). The table summarizes results obtained in the History for Use Case 3-3 Lassa L (dataset 566) and in the History for Use Case 3-3 Ebola (dataset 96). Reconstructed Lassa segment L and Ebola genome sequences are available in Galaxy dataset collections 679 and 115 of their respective histories.

| Virus | BioProject | BioSample id | SRX number | SRR number | Sample ID | BAM file name | Source | Data Type, Selection | Figure, Table from Matranga *et al.* | Metavisitor detection (Trinity) |
| --- | --- | --- | --- | --- | --- | --- | --- | --- | --- | --- |
| EBOV | PRJNA257197 | SAMN03099684 | SRX733660 | SRR1613381 | G3676-2 | G3676-2_S6_L001_001.bam | Human | RNase H | Figure 5 | + |
| EBOV | PRJNA257197 | SAMN03099684 | SRX733656 | SRR1613377 | G3676-2 | G3676-2-std_S13_L001_001.bam | Human | RNA seq | Figure 5 | + |
| EBOV | PRJNA257197 | SAMN03099685 | SRX733661 | SRR1613382 | G3677-1 | G3677-1_S3_L001_001.bam | Human | RNase H | Figure 5 | + |
| EBOV | PRJNA257197 | SAMN03099685 | SRX733657 | SRR1613378 | G3677-1 | G3677-1-std_S10_L001_001.bam | Human | RNA seq | Figure 5 | + |
| EBOV | PRJNA257197 | SAMN03099686 | SRX733662 | SRR1613383 | G3677-2 | G3677-2_S2_L001_001.bam | Human | RNase H | Figure 5 | + |
| EBOV | PRJNA257197 | SAMN03099686 | SRX733658 | SRR1613379 | G3677-2 | G3677-2-std_S9_L001_001.bam | Human | RNA seq | Figure 5 | + |
| EBOV | PRJNA257197 | SAMN03099687 | SRX733663 | SRR1613384 | G3682-1 | G3682-1_S4_L001_001.bam | Human | RNase H | Figure 5 | + |
| EBOV | PRJNA257197 | SAMN03099687 | SRX733659 | SRR1613380 | G3682-1 | G3682-1-std_S11_L001_001.bam | Human | RNA seq | Figure 5 | + |
| LASV | PRJNA254017 | SAMN02927412 | SRX719120 | SRR1595772 | G2431 | LASV678_ERCC117 | Human | RNase H | Figure 2 | + |
| LASV | PRJNA254017 | SAMN02927412 | SRX719079 | SRR1595696 | G2431 | LASV678_ERCC12 | Human | RNA seq | Figure 2 | + |
| LASV | PRJNA254017 | SAMN02927488 | SRX719056 | SRR1595665 | ISTH1003 | LASV347_ERCC126 | Human | RNase H | Figure 2 | + |
| LASV | PRJNA254017 | SAMN02927488 | SRX718926 | SRR1595500 | ISTH1003 | LASV347_ERCC17 | Human | RNA seq | Figure 2 | + |
| LASV | PRJNA254017 | SAMN02927485 | SRX718761 | SRR1594619 | ISTH0531 | LASV334_ERCC136 | Human | RNase H | Figure 2 | + |
| LASV | PRJNA254017 | SAMN02927485 | SRX719205 | SRR1595943 | ISTH0531 | LASV334_ERCC31 | Human | RNA seq | Figure 2 | + |
| LASV | PRJNA254017 | SAMN02927498 | SRX719063 | SRR1595673 | ISTH1121 | LASV363_ERCC69 | Human | RNase H | Figure 2 | + |
| LASV | PRJNA254017 | SAMN02927498 | SRX719134 | SRR1595797 | ISTH1121 | LASV363_ERCC43 | Human | RNA seq | Figure 2 | + |
| LASV | PRJNA254017 | SAMN02927489 | SRX719117 | SRR1595763 | ISTH1038 | LASV349_ERCC62 | Human | RNase H | Figure 2 | + |
| LASV | PRJNA254017 | SAMN02927489 | SRX718979 | SRR1595558 | ISTH1038 | LASV349_ERCC42 | Human | RNA seq | Figure 2 | + |
| LASV | PRJNA254017 | SAMN02927510 | SRX718802 | SRR1594664 | ISTH2050 | LASV386_ERCC84 | Human | RNase H | Figure 2 | + |
| LASV | PRJNA254017 | SAMN02927503 | SRX719192 | SRR1595909 | ISTH2020 | LASV368_ERCC112 | Human | RNase H | Figure 2 | + |
| LASV | PRJNA254017 | SAMN02927484 | SRX718789 | SRR1594651 | ISTH0230 | LASV435_ERCC96 | Human | RNase H | Figure 2 | + |
| LASV | PRJNA254017 | SAMN02927592 | SRX719159 | SRR1595835 | LM032.dep | LM032_Depleted | Mastomys | RNase H | Figure 3 | + |
| LASV | PRJNA254017 | SAMN02927592 | SRX718836 | SRR1594698 | LM032.std | LM032_Standard | Mastomys | RNA seq | Figure 3 | + |
| LASV | PRJNA254017 | SAMN03099734 | SRX733666 | SRR1613388 | NHP_DK9W-AG.dep | 728_Depleted | Macaque | RNase H | Figure 3 | + |
| LASV | PRJNA254017 | SAMN03099735 | SRX733667 | SRR1613389 | NHP_DK9W-AG.std | 728_Standard | Macaque | RNA seq | Figure 3 | + |
| LASV | PRJNA254017 | SAMN03099736 | SRX733668 | SRR1613390 | NHP_DK9W-AL.dep | 729_Depleted | Macaque | RNase H | Figure 3 | + |
| LASV | PRJNA254017 | SAMN03099737 | SRX733669 | SRR1613391 | NHP_DK9W-AL.std | 729_Standard | Macaque | RNA seq | Figure 3 | + |
| LASV | PRJNA254017 | SAMN03099738 | SRX733670 | SRR1613392 | NHP_DK9W-B.dep | 734_Depleted | Macaque | RNase H | Figure 3 | + |
| LASV | PRJNA254017 | SAMN03099739 | SRX733671 | SRR1613393 | NHP_DK9W-B.std | 734_Standard | Macaque | RNA seq | Figure 3 | + |
| LASV | PRJNA254017 | SAMN03099740 | SRX733672 | SRR1613394 | NHP_DK9W-K.dep | 733_Depleted | Macaque | RNase H | Figure 3 | + |
| LASV | PRJNA254017 | SAMN03099741 | SRX733673 | SRR1613395 | NHP_DK9W-K.std | 733_Standard | Macaque | RNA seq | Figure 3 | + |
| LASV | PRJNA254017 | SAMN03099742 | SRX733674 | SRR1613396 | NHP_DK9W-L.dep | 731_Depleted | Macaque | RNase H | Figure 3 | + |
| LASV | PRJNA254017 | SAMN03099743 | SRX733675 | SRR1613397 | NHP_DK9W-L.std | 731_Standard | Macaque | RNA seq | Figure 3 | + |
| LASV | PRJNA254017 | SAMN03099744 | SRX733676 | SRR1613398 | NHP_DK9W-S.dep | 732_Depleted | Macaque | RNase H | Figure 3 | + |
| LASV | PRJNA254017 | SAMN03099745 | SRX733677 | SRR1613399 | NHP_DK9W-S.std | 732_Standard | Macaque | RNA seq | Figure 3 | + |
| LASV | PRJNA254017 | SAMN02927592 | SRX719168 | SRR1595853 | LM032 | LASV68_BLC | Mastomys | RNA seq | Figure 4, Table 1 | + |
| LASV | PRJNA254017 | SAMN02927476 | SRX727329 | SRR1606288 | G733 | LASV_90 | Human | RNA seq | Figure 4, Table 1 | + |
| LASV | PRJNA254017 | SAMN02927592 | SRX733690 | SRR1613412 | LM032 | LM032_HS | Mastomys | Hybrid Selection | Figure 4, Table 1 | + |
| LASV | PRJNA254017 | SAMN02927476 | SRX733681 | SRR1613403 | G733 | G733_HS | Human | Hybrid Selection | Figure 4, Table 1 | + |
| LASV | PRJNA254017 | SAMN02927593 | SRX727318 | SRR1606277 | LM222 | LASV_74 | Mastomys | RNA seq | Table 1 | + |
| LASV | PRJNA254017 | SAMN03099732 | SRX733664 | SRR1613386 | Z002 | LASV_77 | Mastomys | RNA seq | Table 1 | - |
| LASV | PRJNA254017 | SAMN03099733 | SRX733665 | SRR1613387 | G090 | LASV_79 | Human | RNA seq | Table 1 | + |
| LASV | PRJNA254017 | SAMN02927477 | SRX727310 | SRR1606267 | G771 | LASV94 | Human | RNA seq | Table 1 | + |
| LASV | PRJNA254017 | SAMN02927399 | SRX734464 | SRR1614275 | G2230 | Solexa-100929.tagged_332 | Human | RNA seq | Table 1 | + |
| LASV | PRJNA254017 | SAMN02927483 | SRX731079 | SRR1610580 | ISTH0073 | Solexa-106870.tagged_851 | Human | RNA seq | Table 1 | + |
| LASV | PRJNA254017 | SAMN02927500 | SRX719163 | SRR1595846 | ISTH1137 | LASV353_BLC | Human | RNA seq | Table 1 | + |
| LASV | PRJNA254017 | SAMN02927503 | SRX718749 | SRR1594606 | ISTH2020 | LASV368_ERCC03 | Human | RNA seq | Table 1 | + |
| LASV | PRJNA254017 | SAMN02927504 | SRX727274 | SRR1606236 | ISTH2025 | LASV374_ERCC58 | Human | RNA seq | Table 1 | + |
| LASV | PRJNA254017 | SAMN02927510 | SRX718860 | SRR1594723 | ISTH2050 | LASV386_ERCC48 | Human | RNA seq | Table 1 | + |
| LASV | PRJNA254017 | SAMN02927484 | SRX718809 | SRR1594671 | ISTH0230 | LASV435_ERCC53 | Human | RNA seq | Table 1 | + |
| LASV | PRJNA254017 | SAMN02927593 | SRX733692 | SRR1613414 | LM222 | LM222_HS | Mastomys | Hybrid Selection | Table 1 | + |
| LASV | PRJNA254017 | SAMN03099732 | SRX733678 | SRR1613400 | Z002 | Z002_HS | Mastomys | Hybrid Selection | Table 1 | + |
| LASV | PRJNA254017 | SAMN03099733 | SRX733679 | SRR1613401 | G090 | G090_HS | Human | Hybrid Selection | Table 1 | + |
| LASV | PRJNA254017 | SAMN02927477 | SRX733682 | SRR1613404 | G771 | G771_HS | Human | Hybrid Selection | Table 1 | + |
| LASV | PRJNA254017 | SAMN02927399 | SRX733680 | SRR1613402 | G2230 | G2230_HS | Human | Hybrid Selection | Table 1 | + |
| LASV | PRJNA254017 | SAMN02927483 | SRX733683 | SRR1613405 | ISTH0073 | ISTH0073_HS | Human | Hybrid Selection | Table 1 | + |
| LASV | PRJNA254017 | SAMN02927500 | SRX733685 | SRR1613407 | ISTH1137 | ISTH1137_HS | Human | Hybrid Selection | Table 1 | + |
| LASV | PRJNA254017 | SAMN02927503 | SRX733686 | SRR1613408 | ISTH2020 | ISTH2020_HS | Human | Hybrid Selection | Table 1 | + |
| LASV | PRJNA254017 | SAMN02927504 | SRX733687 | SRR1613409 | ISTH2025 | ISTH2025_HS | Human | Hybrid Selection | Table 1 | + |
| LASV | PRJNA254017 | SAMN02927510 | SRX733688 | SRR1613410 | ISTH2050 | ISTH2050_HS | Human | Hybrid Selection | Table 1 | + |
| LASV | PRJNA254017 | SAMN02927484 | SRX733684 | SRR1613406 | ISTH0230 | ISTH0230_HS | Human | Hybrid Selection | Table 1 | + |
| LASV | PRJNA254017 | SAMN02927592 | SRX733689 | SRR1613411 | LM032 | LM032_Depleted | Mastomys | cDNA | ND, manually added to the original sup file 3 | + |
| LASV | PRJNA254017 | SAMN02927592 | SRX733691 | SRR1613413 | LM032 | LM032_Standard | Mastomys | cDNA | ND, manually added to the original sup file 3 | - |

The results of both analyses are summarized in Table 4. Metavisitor was able to detect Ebola virus in all corresponding sequence datasets as well as Lassa virus in 53 out the 55 sequence datasets generated from Lassa virus samples. Importantly, Matranga et al also did not report reconstructed Lassa genomic segments from the two datasets that we found to be Lassa virus negative, likely due to high read duplication levels in the corresponding libraries [28]. The reconstructed Lassa virus L segments are compiled in the dataset 679 of History for Use Case 3-3 Lassa L whereas reconstructed Ebola virus genomes are compiled in the dataset 115 of History for Use Case 3-3 Ebola. In these sequences, *de novo* assembled segments in uppercases are integrated in the reference guide sequence (lowercase) used for the reconstruction. To note, for viruses with segmented genomes the workflow for Use Case 3-3 has to be used separately with corresponding guide sequences for the appropriate segment to be reconstructed. As an example, we used this workflow with the input variables “Lassa” (filter term for the “Parse blast output and compile hits” tool) and “NC_004296.1” (Lassa S segment used for guiding the reconstruction) to generate the History for Use Case 3-3 Lassa S.

At this stage, users can use the genomic fasta sequences for further analyses. For instance, multiple sequence alignments) can be directly performed for phylogenetics or variant analyses, or reads in the original datasets can be realigned to the viral genomes as in Use Cases 1 and 2 to visualize their coverages.

## Discussion

In order to address accessibility, reproducibility and transparency issues in bioinformatics analyses for the detection and reconstruction of viruses, we developed Metavisitor, an open-source suite of tools and workflows executable in Galaxy. Galaxy provides a framework supported by a growing community, and allows executing computational tools and workflows through a user-friendly web interface without need for advanced skills in bioinformatics. Thus, on the one hand, Metavisitor may be useful to many researchers, from seasoned bioinformaticians to medical virologists trying to identify the source of an unknown illness. On the other hand, the advanced Galaxy functionalities ensure the highest levels of computational analyses, through rigorous recording of the produced data and metadata and of the used parameters as well as the ability to share, publish and reproduce these analyses, as illustrated by this work. Another major benefit from their integration in Galaxy is that, as any Galaxy workflow, the Metavisitor workflows may easily be adapted, modified or extended with tools from the active Galaxy developer community.

Through use cases, we have shown that the current set of Metavisitor tools can generate workflows adapted to diverse situations: (a) Short or longer reads from small RNAseq, RNAseq or DNAseq can be used as input data, in a fastq or fasta format, with or without clipping adapter sequences. (b) Sequence information in these input data can be used as is, or compressed using our reads-to-sequences procedure or normalization by median [24]. This compression greatly reduces the workload and may improve the quality of the *de novo* assembly step (see Use Cases 1-1 to 1-3). (c) We used three alignment tools in this work based on Bowtie or Bowtie2, including our sRbowtie tool adapted to short RNA reads. Indeed any alignment software producing BAM/SAM outputs may be used in future Metavisitor workflows, provided that they are wrapped for their integration in the Galaxy framework. (d) We have shown the benefit of subtracting non-viral reads before *de novo* assembly by their alignment to host, parasite or symbiont genomes. Nevertheless, Use Case 3-3 illustrates that this step is optional when experimental procedures generate sequence datasets highly enriched in viral sequences. (e) We adapted efficient workflows for two *de novo* assembler programs (Oases and Trinity). It is noteworthy that both of these assemblers could be used in parallel in a single Metavisitor workflow to produce more contigs, which are subsequently filtered by blastn/x alignments to known viral sequences. Any other *de novo* assembly software can be adapted to be used as Metavisitor assembly tool. (f) The viral genome reconstruction can also be adapted. We found that when the number of blast hits to the guide sequences is high, indicative of a high coverage, then the CAP3 assembly of the corresponding hit sequences may be omitted. For instance, our tool “blast_to_scaffold” was sufficient to generate full Lassa and Ebola genome reconstructions in Use Case 3-3. (g) Finally, central to Metavisitor are the viral nucleotide and protein references used in the workflows to identify viral contigs or viral reads. We retrieved the vir1 references from the NCBI using the “Retrieve FASTA from NCBI” Galaxy tool with an explicit query string. We will re-run this tool on a regular basis with the same query string to update the vir1 references with sequences newly deposited to the NCBI databases. However, users are free to adapt Metavisitor workflows to their own viral references either by running the “Retrieve FASTA from NCBI” tool using query strings of there choice or by uploading their own fasta sequences. This is possible since alignment tools in the Metavisitor tool suite, including sRbowtie, Bowtie, Bowtie2 and Blast, can work with indexes generated on the fly from fasta datasets present in Galaxy users’ histories.

### A modular and scalable software for biologists and clinicians

Metavisitor tools are modules that can be combined by biologists and clinicians to build analyses workflows adapted to their needs: from detection or reconstruction of known viruses in Drosophila small RNA-seq datasets to novel virus discovery in Anopheles [25] to diagnosis and reconstruction of viruses of patients from RNA-seq datasets.Importantly we showed that Metavisitor is able to detect co-infections by multiple viruses(see Use Case 1-4 for an example).

Viral genome sequences reconstructed by Metavisitor can be used for any subsequent analysis, including phylogenetic or genetic drift analyses in contexts of epidemics or viruses surveillance in field insect vectors, animal or human populations, and systematic identification of viruses for evaluation of their morbidity. The use of Galaxy dataset collections allows to adapt Metavisitor to high throughput analyses. For instance, in Use Cases 3-1 to 3-3 we were able to analyze in batch dozens of patient data from multiplexed sequencing experiments, with consistent tracking of individual samples, from fastq datasets to individual viral genome reconstruction. Thus we are confident that Metavisitor is scalable to large epidemiological studies or to clinical diagnosis in hospital environments. One possible immediate exploitation of this scalability would be to reconstruct sequences of the Zika virus strains from infected patients and identify possible co-infections that could explain and correlate with clinical symptoms.

### Future directions

The central idea in Metavisitor is to detect *de novo* contigs of viral sequence reads through blast alignments. Indeed, the ability to form *de novo* viral contigs that align to the large viral sequence database extracted from Genbank NCBI (vir1) provides very strong evidence of the presence of a virus while ensuring a low rate of false positives. However, this current vir1 reference database is redundant and contains sequences whose annotations are misleading or not meaningful (for instance, chimeric sequences between host and viral genomes or patent sequences). We will work at removing redundant or mis-annotated sequences from vir1. This curated vir1 reference will improve the speed of the alignment steps, reduce the size of the reports generated by Metavisitor while including phylogenetic informations on detected viruses. In the meantime, the versatility of Metavisitor allows users to work with their own viral sequence references.

We are aware that low viral loads in sequenced samples and/or viral read alignments to scattered short regions of viral genomes may result in failure to assemble viral contigs and thus in putative false negatives. We have shown how to keep track of these false negatives in the Workflow for Use Case 3-2 by parsing the SAM alignments to the vir1 index in order to annotate and to count viral reads before the contig assembly steps (Table 3). This tracking of putative false negatives will also be simplified with a curated vir1 reference.

We finally wish to stress that Metavisitor has the potential for integrating detection or diagnosis of non-viral, microbial components in biological samples. Eukaryotic parasites or symbionts and bacteria are mostly detectable in sequencing datasets from their abundant ribosomal RNAs whose sequences are strongly conserved in the main kingdoms. This raises specific issues for their accurate identification and their taxonomic resolution which are not currently addressed by Metavisitor. However, many tools and databases [29] addressing these metagenomics challenges can be adapted, when not already, to the Galaxy framework. For instance, Qiime [30] and the SILVA database of ribosomal RNAs [31] can be used within Galaxy and could thus be integrated in future Metavisitor workflows.

## Funding

This work received financial support to CA from the Agence Nationale de la Recherche grant ANR-13-BSV2-0007 « Plastisipi » and from the Institut Universitaire d’Ingéniérieen Santé IUIS grant “ARTiMED” and to KDV from the European Research Council (http://erc.europa.eu/funding-and-grants), Support for frontier research, Advanced Grant #323173 AnoPath; European Commission (http://ec.europa.eu/research/fp7), FP7 Infrastructures #228421 Infravec; and French Laboratoire d’Excellence (http://www.enseignementsup-recherche.gouv.fr/cid55551/investissements-d-avenir-projets-laboratoires-d-excellence-par-region-et-domaine.html), “Integrative Biology of Emerging Infectious Diseases” #ANR-10-LABX-62-IBEID.

The funders had no role in study design, data collection and analysis, decision to publish,or preparation of the manuscript.

## Acknowledgements

We thank the Galaxy community for their support, Eugeni Belda and Emmanuel Bischoff for helpful discussions and Julie Reveillaud for critical reading of the manuscript. GC, MvdB and CA conceived the project. CA and MvdB developed and implemented tools in the Galaxy framework. GC, MvdB, JP and CA performed bioinformatics analysis. GC, MvdB, KV and CA wrote the manuscript. CA and KV provided funding. All authors read and approved the final manuscript.

## Supplementary Information

### Supplementary File 1

MAFFT (http://www.ebi.ac.uk/Tools/msa/mafft/) Multiple Alignment of the Nora virus genome sequences published (JX220408.1 and NC_007919.3) or generated in Use Cases 11 to 13 (Nora_MV, Nora_raw_reads and Nora_Median-Norm-reads). A view of the alignments was produced by MView (http://www.ebi.ac.uk/Tools/msa/mview/). The html file can be visualized by opening it locally in any web browser software.

**Supplementary Figure 1.**
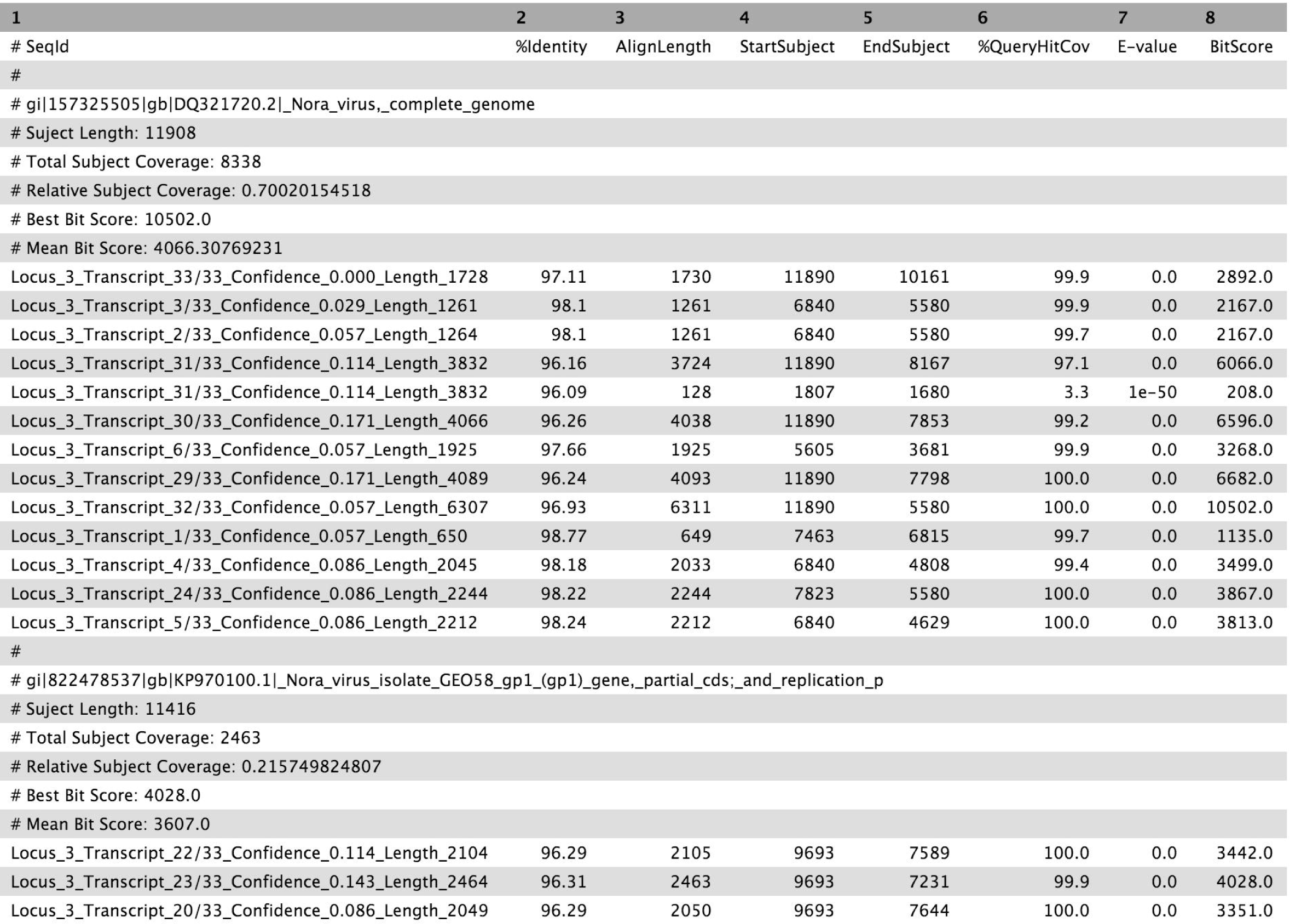
Screenshot of an output produced by the “Parse blast output and compile hits” Metavisitor tool.

